# Subclonal architecture, evolutionary trajectories and patterns of inheritance of germline variants in pediatric glioblastoma

**DOI:** 10.1101/434241

**Authors:** Mary Hoffman, Aaron H Gillmor, Daniel J Kunz, Michael Johnston, Ana Nikolic, Kiran Narta, Jennifer King, Katrina Ellestad, Ngoc Ha Dang, Florence MG Cavalli, Michelle M Kushida, Fiona J Coutinho, Betty Luu, Yussanne Ma, Andrew J Mungall, Richard Moore, Marco A Marra, Michael D Taylor, Trevor J Pugh, Peter B Dirks, Donna L Senger, Benjamin D Simons, Jennifer A Chan, A Sorana Morrissy, Marco Gallo

## Abstract

Pediatric glioblastoma (pGBM) is a lethal cancer with no effective therapies. Intratumoral genetic heterogeneity and mode of tumor evolution have not been systematically addressed for this cancer. Whole-genome sequencing of germline-tumor pairs showed that pGBM is characterized by intratumoral genetic heterogeneity and consequent subclonal architecture. We found that pGBM undergoes extreme evolutionary trajectories, with primary and recurrent tumors having different subclonal compositions. Analysis of variant allele frequencies supported a model of tumor growth involving slow-cycling cancer stem cells that give rise to fast-proliferating progenitor-like cells and to non-dividing cells. pGBM patients’ germlines had subclonal structural variants, some of which underwent dynamic frequency fluctuations during tumor evolution. By sequencing germlines of mother-father-patient trios, we found that inheritance of deleterious germline variants from healthy parents cooperate with *de novo* germline and somatic events to the tumorigenic process. Our studies therefore challenge the current notion that pGBM is a relatively homogeneous molecular entity.

---

Pediatric glioblastoma (pGBM) is a lethal brain tumor. Unlike adult GBM, no effective therapeutic intervention or standard of care currently exists for this cancer. Despite their histological similarities, pGBM is clinically, biologically and molecularly distinct from adult GBM *(1)*. Unfortunately, the biological principles uncovered by studying the adult disease have not translated to the pediatric malignancy *(2)*. A major breakthrough in understanding the molecular principles underlying pGBM stemmed from the discovery that a third of patients harbor somatic mutations in *H3F3A*, a gene encoding the histone variant H3.3 *(3, 4)*. Mutations in this gene cause one of two amino acid substitutions in H3.3: (i) Mutation of lysine 27 to methionine (K27M); (ii) mutation of glycine 34 to arginine (G34R) or valine (G34V). Both types of mutations interfere with the wildtype function of H3.3 *(5, 6)*, but the H3.3K27M variant has been studied more extensively because it is more frequent and trimethylation of K27 has an established and important role in repressing gene transcription and compacting chromatin. Several experimental models have been used to show that *H3F3A* mutations can induce cellular hyperproliferation on their own, but they need to cooperate with other drivers, including *TP53* mutations, to produce overt malignancies *(6–8)*. For the majority of pGBM patients without H3.3 mutations, the early genetic events leading to overt malignancy are not fully understood.

It is currently believed that pGBM, similarly to other pediatric cancers, is characterized by relatively bland genomes *(9)*. Furthermore, *H3F3A* mutations were shown to be clonal in pGBM *(10)*, and to result in global loss of trimethylation of lysine 27 of histone 3 (H3K27me3) in virtually all cells in a tumor *(6)*. These findings have contributed to the view that pGBM might have a relatively low level of intratumoral complexity at the genomic and functional level. However, a recent report identified differences in mutational signatures between matched primary and recurrent pGBM samples using exome sequencing *(10)*.

Evidence of subclonal architecture in diffuse intrinsic pontine glioma and pGBM was recently provided *(11)*. The authors reanalyzed previously published exome datasets for 36 pGBM samples and identified evidence for the coexistence of subclones in each tumor. To the best of our knowledge, no study to date has systematically assessed subclonal composition and evolution in matched primary and recurrent samples from the same patient using whole-genome information. Such studies are important because they may allow inference of biological principles responsible for tumor evolution and recurrence, and they might identify intergenic variants with important roles in the etiology of this cancer. Furthermore, treatment regimens for pGBM differ between centers and individual oncologists, raising the possibility that different therapeutic approaches might skew recurrences toward the acquisition of specific molecular features. The relative rarity of pGBM (adjusted incidence rate of 0.15/0.16 in 100,000 population) *(12)* and the lack of standardized practices for surgical resection contribute to the scarce availability of diagnostic-recurrent sample pairs for study.

New whole-genome technologies that allow sensitive measurements of cellular heterogeneity in a single tumor, and documentation of tumor evolution over time, have yet to be systematically employed for this malignancy. We hypothesized that whole-genome sequencing (WGS) of primary and recurrent tumors, coupled with targeted bioinformatic approaches to specifically test for subclonal architecture of each tumor, might bring new insight into the intratumoral organization of pGBM and its evolution. Understanding the principles exploited by the tumor to escape treatment may have important implications for rationally selecting or designing personalized treatment options for these rare and aggressive cancers.

## Results

### pGBM samples and whole-genome sequencing with linked-reads

We characterized longitudinally collected pGBM samples that include primary tumors and their recurrences, in addition to peripheral blood (used as germline control), and were biobanked at our institution (Calgary cohort). For 4 out of 5 patients we had at least two longitudinal samples and for one patient we had three recurrent samples (**Table S1**). For the purposes of this manuscript, diagnostic samples are referred to as “sample 1” and each consecutive resection from the same patient is referred to as “sample *n*”. Treatment history for each patient is known (**Table S2**). We also obtained a set of matched germline and primary pGBM samples biobanked in a second institution (Toronto cohort) (**Table S3**) and used these for validation purposes. We extracted high molecular weight DNA (average length distribution > 40 kbp) from each sample and generated libraries for WGS with linked-read technology (10xGenomics). With this system, short next-generation sequencing reads originating from the same DNA molecule (ie an individual chromosome fragment) are tagged with an identical barcode, enabling the assembly of Mb-size haplotypes from individual chromosomes. Average sequencing depth for all samples was 35.6x, while average physical coverage was 178x because of the use of linked reads (**Table S4**). The longest phase block generated was 71 Mbp, with an average maximum length for all samples of 16.6 Mbp and a median of 12 Mbp (**Table S4**).

### Diagnostic and recurrent pGBM samples share few single nucleotide variants

We used linked-read WGS data for pGBM samples and matched germlines to call somatic variants in diagnostic samples and their recurrences. Focusing specifically on non-synonymous single-nucleotide variants (SNVs), our data are in agreement with previous reports of low mutation burden *(3, 13, 14)* in pGBM (range: 26-124 SNVs per tumor; average: 65.6 SNVs per tumor in primary samples, 60 SNVs per tumor in the recurrences). Overall, we found that recurrent tumors shared few somatic mutations with their matched primary malignancies, with only between 1 and 5 non-synonymous SNVs shared between longitudinal samples from the same patient. This translated to an average of 7.8% non-synonymous SNVs that were shared between a primary tumor and its recurrence. A similar amount (11.8%) of somatic SNVs and indels were shared between diagnostic and recurrent samples in the pediatric brain tumor medulloblastoma *(15)*, in contrast with the much higher proportion shared between primary and recurrent adult glioma samples (54%) *(16)*. Surprisingly, we observed that the total number of somatic SNVs tended to be stable in recurrences post-treatment (**Fig. 1A-1E**; **Supplemental file 1**), whereas mutational load is known to increase 5-fold at recurrence in pediatric medulloblastoma *(15)* as well as in adult gliomas *(16)*. Patient 1 was one exception (**Fig. 1A**), but this individual harbored a germline deletion of the last exon of *TP53* (**Fig. S1A and S1B**), which might have contributed to accelerated tumor evolution by increasing mutational events. Sample 2 had 126 somatic non-synonymous SNVs, and only lesions in *ARMC9* and *DSE* were shared with the diagnostic sample. Interestingly, *DSE* encodes a dermatan sulfarase epimerase and is expressed in the human brain at higher levels before birth than after birth (**Fig. S1C and S1D**). Patient 2 had a somatic mutation in *H3F3A* that was retained in the second sample (**Fig. 1B**), together with a mutation in *MUC22* and two deleterious SNVs in *TP53*. This patient was treated with temozolomide and local radiation (**Supplementary Table S2**), which is the standard of care for adult GBM patients. However, the overall number of somatic, non-synonymous SNVs was lower in the second sample (69 SNVs) than in the diagnostic sample (109 SNVs), whereas this treatment is known to increase mutational burden in adults. This difference in response to chemotherapy and radiation between pGBM and adult GBM was also noted in an independent cohort of patients in a recent publication *(10)*. Samples 1 and 2 from patient 3 only shared a mutation in *PCNX* (**Fig. 1C**). Patient 4 had 84 somatic non-synonymous SNVs in the diagnostic sample (**Fig. 1D**), and no recurrence was available for sequencing. Patient 5 had 26 non-synonymous SNVs in the diagnostic sample and in sample 2, 38 in sample 3, and 39 in sample 4 (**Fig. 1E**). SNVs in 5 genes were shared between sample 1, sample 2 and sample 3 (*ELAVL2*, *VWF*, *KCNJ12*, *HEXIM2* and *ZNF816*). Only SNVs in *VWF* (encoding the von Willebrand factor) and *KCNJ12* (encoding an ion channel) were conserved between all samples for this patient.

**Fig. 1.**
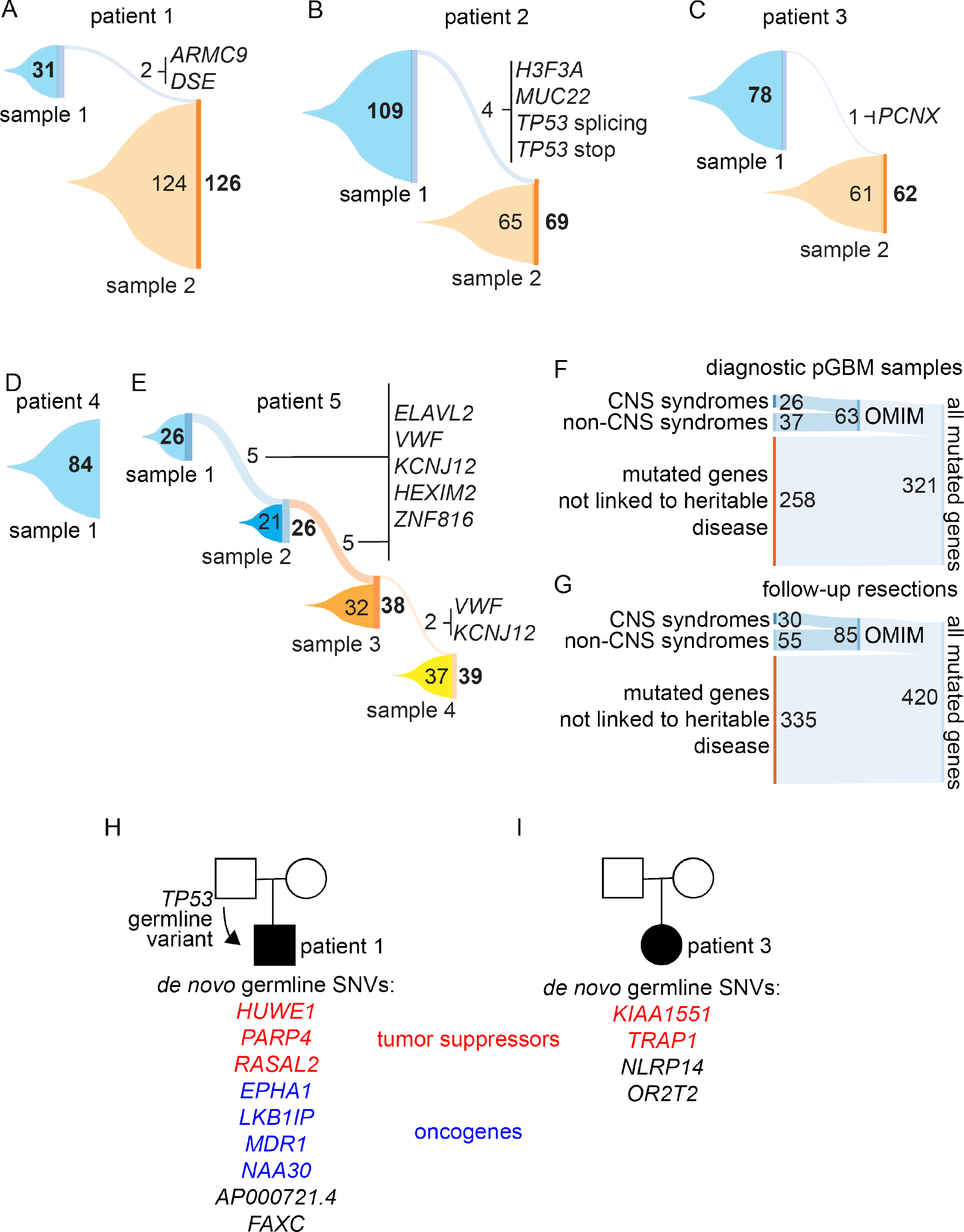
Single nucleotide variants in pediatric GBM samples. Sankey plots for patients 1 (A), 2 (B), 3 (C), 4 (D) and 5 (E). Each plot illustrates changes in the mutational landscapes between primary tumors and recurrences. Bold numbers represent total number of mutations in each profiled tumor. Gene lists represent the mutated genes that are shared between consecutive tumors. Analysis of all exonic, non-synonymous single nucleotide variants (SNVs) in all primary tumors reveals a fraction of genes are associated with hereditary syndromes, according to the OMIM database. Out of the 63 genes with links to hereditary disease, 26 were specifically linked to central nervous system (CNS) syndromes. Analysis of all exonic, non-synonymous SNVs in all recurrent tumors. Out of 85 genes with links to hereditary disease, 30 were specifically linked to CNS syndromes. Germlines were sequenced for patient 1, and the patient’s mother and father. A deletion of the last exon of *TP53* was detected in the father, and it was inherited by patient 1. Non-synonymous SNVs in 9 other genes were identified in patient’s 1 germline, but not in the parents. (F) *De novo* non-synonymous SNVs in four genes were identified in patient’s 3 germline.

No SNV was shared among all samples. Only 4 genes were mutated in at least two primary tumors (**Table S5**), including *VWF*. Six genes were mutated in at least two recurrences from different patients (**Table S6**). No established oncogene or tumor suppressor was recurrently mutated in our cohort, with the exception of *TP53* (altered in two patients). Our data therefore show that pGBM is characterized by large intertumoral genetic heterogeneity and by large evolutionary changes following surgical resection and treatment.

No obvious pattern of gene mutations was observed in our dataset. Gene ontology analysis did not identify any significantly enriched class of mutated genes. However, we found that non-synonymous SNVs in diagnostic samples were significantly enriched for genes with an OMIM entry and linked to hereditary syndromes (19.6% of the 321 mutated genes, permutation test p = 0.00082; **Fig. 1F**, **Fig. S1E,F**). 41.3% of these genes are associated with syndromes of the central nervous system (CNS). Similarly, non-synonymous SNVs in subsequent resections had a significant enrichment for genes with OMIM entries (20.2% of the 420 mutated genes, permutation test p < 10^−5^; **Fig. S1E,F**), and 35.2% of these genes were associated with hereditary CNS syndromes (**Fig. 1G**). These observations suggest that disruption of multiple unrelated networks of genes may be sufficient for pGBM initiation and progression. The identification of deleterious SNVs in genes linked to hereditary syndromes is intriguing, given the early-life onset of this malignancy.

### Germline sequencing of mother-father-patient trios

Intrigued by the identification of somatic non-synonymous SNVs in genes associated with hereditary disorders, we asked whether pGBM behaves as a developmental syndrome. We reasoned that if we assumed cancer was a developmental disorder of progressive mutagenesis, individual patients might undergo systemic accumulation of mutations over time. We decided to investigate this possibility by assembling trios composed of mother, father and patient, implementing an approach similar to what is done with genetic disorders. We were able to obtain the germlines (peripheral blood) of the parents of patient 1 and patient 3, and used them for WGS with linked reads. Analysis of these trios enabled us to categorize non-synonymous SNVs as either unique to the pGBM patient germline, or inherited from one of the parental genomes. Our data show that a *TP53* germline variant identified in patient 1 (**Fig. S1A and S1B**) was inherited from one of the parents. In addition, non-synonymous SNVs in 9 genes were identified uniquely in the germline of this patient (**Fig. 1H**). Among these genes, *RASAL2* encodes a RasGAP and is a tumor suppressor *(17)*; *PARP4* encodes a member of the poly(ADP)-ribose family, and was recently shown to be mutated in the germlines of patients with thyroid and breast cancer *(18)*; *HUWE1* encodes an E3 ubiquitin ligase that was shown to be a tumor suppressor in mouse models of skin cancer *(19)* and colorectal cancer *(20)*. Similarly, we called 4 *de novo* non-synonymous SNVs in the germline of patient 3 (**Fig. 1I**). Among these genes, *KIAA1551* was recently identified as a candidate tumor suppressor and is frequently deleted in different cancer types *(21)*. Overall, we noticed a significant increase in the number of short (30-50,000 bp) deletions in the germlines of pGBM patients (n = 8) compared to the control (parental) germlines (n = 4) we sequenced (Mann Whitney test p = 0.0040; **Fig. S1G**). This collection of sequenced trios is a unique dataset in the context of pGBM and offers a new perspective on the emergence of germline variants in pediatric patients with sporadic cancer.

### Subclonal architecture of pGBM at diagnosis and recurrence

We next asked whether the small proportion of SNVs shared between each diagnostic sample and its recurrence were a reflection of unexpected intratumoral genomic complexity and consequent clonal selection. We assessed the presence of genetically distinct subclonal cell lineages in each tumor before and after treatment by using somatic mutations and copy number alterations to build a genetic phylogeny for each patient using the EXPANDS algorithm *(22)*. These phylogenies show that each pGBM and its recurrence(s) are characterized by extensive intratumoral genomic heterogeneity and complex subclonal architecture (**Fig. 2A-E**). For instance, in the diagnostic sample from patient 1 we detected 8 subclonal lineages that correspond to three main groups or clusters (**Fig. 2A**). Subclones e and f (1ef) represent ancestral genetic lineages not far diverged from the germline, while subclones 1abcdg and 1h form two major phylogenetic branches, exhibiting moderate to extreme divergence from the germline. In all patients and all samples we observe distinct phylogenetic branches. These support the presence of multiple genetically distinct subclones that change significantly in frequency throughout the evolution of a tumor. Importantly, each recurrent tumor also had a few subclones clustering close to the germline, likely indicating the stable presence of early (ancestral) clones which are not actively accumulating somatic alterations. This observation is best illustrated in patient 5 where three recurrences were available (**Fig. 2E**). Throughout this patient’s tumor progressive divergence from the germline occurs at each successive recurrence in a number of genetic lineages, yet a number of ancestral subclones remain detectable at all timepoints (e.g. 2agh at the first recurrence; 3f at the second recurrence; 4ef at the third recurrence). While these subclones likely had ancestral roles in the early part of the evolutionary path of this patient’s tumor, each subsequent recurrence is marked by distinct lineages that have diverged significantly from this baseline.

**Fig 2.**
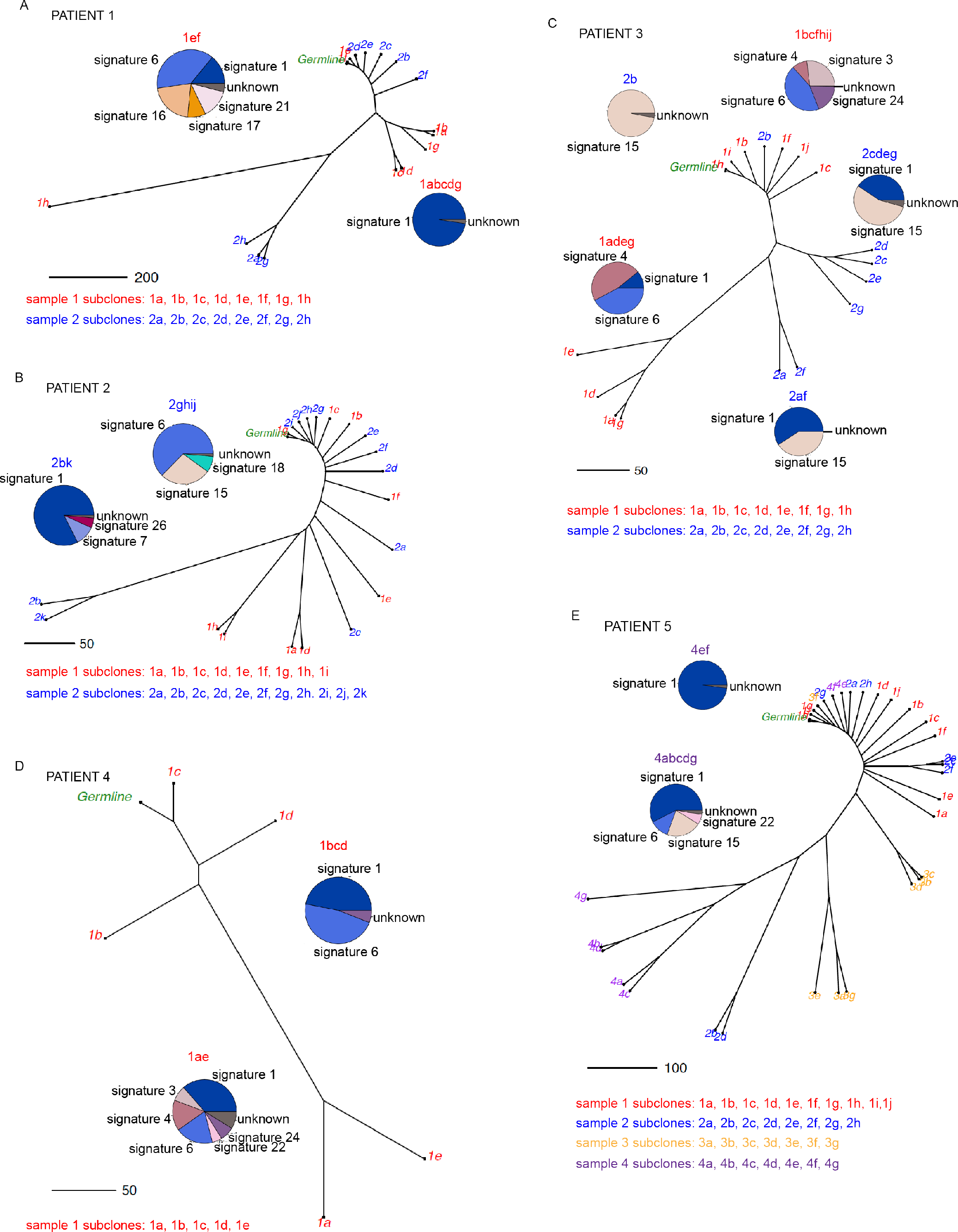
Subclonal architecture and evolution in pediatric GBM. Phylogenetic trees reflect intermixing of genetic lineages in each sample from patients 1 (A), 2 (B), 3 (C), 4 (D), and 5 (E). The subclonal structure of each tumor sample was inferred using somatic mutations and copy number alterations. Distance between nodes is proportional to the number of genetic differences between subclones. Branch tips are color-coded according to sample number, and labeled according to biopsy number (1, 2, 3) and clonal lineage (a, b, c). For selected clusters of genetic subclones, pie charts display the contribution of mutational signatures detectable using the somatic mutations specific to those phylogenetic branches.

In addition to clonal composition, somatic SNVs were further used to infer mutational signatures. Having already inferred the phylogenetic lineages present in each sample let us select SNVs corresponding to distinct clusters of subclones, such that we could then compare the range of mutation signatures from subclones within each sample. In each patient we see evidence for multiple mutation signatures, indicating the influence of more than one mutagenic process. For instance, in the diagnostic sample of patient 2, the subclones clustering near the germline (1bcfhij) were characterized by mutational signatures 3,4,6, and 24, while the more evolutionarily divergent subclones accumulated mutations dominated by signatures 4, 6 and 1 (**Fig. 2C**). Similar patterns of divergence between mutational signatures specific to clusters of subclones were observed for all other patients (**Fig. 2B-E** and **Fig. S2**). Signature 1 results from the spontaneous deamination of 5-methylcytosine, is the most commonly seen across cancer types, and was observed in our cohort in all samples though at different levels in each subclone (**Fig. S2**). Signatures 6 and 15, prevalent in cancers with defective mismatch repair, were frequently observed in many subclones, with a handful of additional signatures (including 8; previously observed in the pediatric brain tumor medulloblastoma) detectable in individual subclones. All together, these data indicate that different mutational processes may operate at different times during the evolution of individual tumors. Alternatively, different mutational processes are acting concurrently at the onset, and the changing selective pressure present at each treatment stage enables different genetic lineages to become dominant within the tumor.

### Identification of structural variants in pGBM

pGBM is known to harbor somatic structural variants (SVs), some of which recur at low frequency in the patient population (*23, 24*). Because WGS with linked reads data are optimal for identification and visualization of structural variants and subclonal events, we explored the behavior of SVs during tumor evolution. SVs were called with the longranger package. We defined large SVs as variants affecting > 30 kbp of the genome. We processed our linked-read WGS data for germlines and tumors independently of each other. This approach allowed us to retain SVs that might be important for tumor etiology, but that would otherwise have been filtered out because they are also present in the germlines. With this approach, we identified 20 to 30 SVs in the germlines of each pGBM patient compared to the reference genome (**Fig. 3A-E**; **Supplemental file 2**). The overall number of large SVs did not change dramatically between the primary and recurrent tumors. Patient 1 was the only one displaying a large increase in the number of SVs in the primary tumor compared to the germline (**Fig. 3A**). This was probably because of chromosome 6 chromothripsis events, which represented 62% of total SVs for this patient. No trace of chromosome 6 chromothripsis was observed in the second resection and the total number of SVs was similar to the germline. This was unexpected because most patients were treated with radiation, which is known to induce SVs. However, this absence is in accord with the observed divergent clonal structure of pGBM (**Fig. 2**), and suggests that the chromothriptic event may have been present in a subclone. Adult gliomas experience a large number of mutational events (SVs and SNVs) during standard of care treatment (radiation plus temozolomide) *(16)*. Patient 2 underwent radiation and temozolomide treatment, yet the overall mutational burden did not drastically increase (**Fig. 3B**). We detected SVs affecting previously known cancer genes as well. Sequencing data provide evidence of a duplication of *NTRK2*, the overexpression of which was previously shown to contribute to tumor metastasis *(25)*. Gene fusions involving *NTRK2* have been previously reported in pGBM *(24)*. Patient 3 had a somatic *ETV6-NTRK3* gene fusion in both primary and recurrent tumors. This gene fusion has been reported in pGBM before *(24)*, and it was also shown to be have oncogenic function in other malignancies, including leukemia *(26)*, breast cancer *(27)* and papillary thyroid carcinoma *(28)*. We also identified a translocation predicted to truncate the proto-oncogene *MET* in patient 4. *MET* amplifications are relatively common in adult GBM *(29)*, and lesions in this gene are found in several cancer types *(30)*, including pGBM *(24, 31)*.

**Fig. 3.**
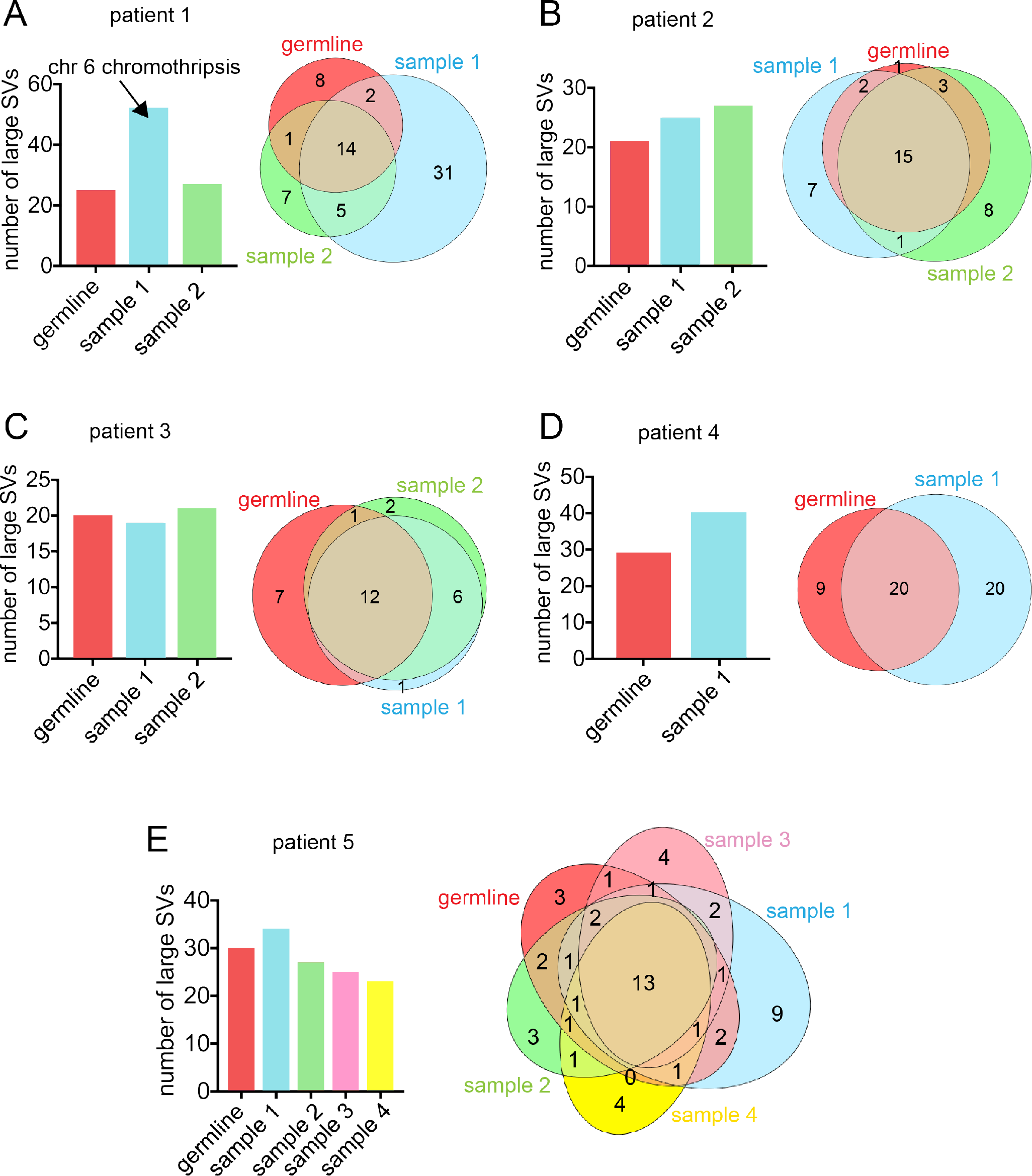
Number of large structural variants (SVs) in the germlines and pGBM tumor tissue. Total number of large SVs in the germline, primary and recurrent tumors for patients 1 (A), 2 (B), 3 (C), 4 (D), and 5 (E). SV calls were performed with the LongRanger pipeline.

We have identified several SVs that recurred in more than one patient in our cohorts. Among these, we found germline deletions affecting the coding sequences of *BTNL3*, *BTNL8*, *APOBEC3A* and *APOBEC3B*. Interestingly, *BTNL8* encodes a cell surface protein involved in T cell activation *(32)*, whereas the *APOBEC3* genes encode cytidine deaminases with roles in antiviral immunity *(33)*. These deletions abrogated gene expression (**Fig. S3**).

### Germline and somatic structural variants during tumor evolution

In some instances, tumors had lower numbers of SVs than their matched germlines (see diagnostic sample in **Fig. 3C** and samples 2-4 in **Fig. 3E**). We found that every patient carried some germline SVs that were not detected in their diagnostic samples and their recurrences. There were 8 such germline-specific SVs for patient 1 (**Fig. 3A**), 1 for patient 2 (**Fig. 3B**), 7 for patient 3 (**Fig. 3C**), 9 for patient 4 (**Fig. 3D**) and 3 for patient 5 (**Fig. 3E**). Each SV call was made with longranger and was manually reviewed, checked for coverage in the region of the call and confirmed. Some SVs with allelic frequencies between 0.4 and 0.7 in the germline also underwent fluctuations in the tumors, while others were stable (**Fig. S4A-D**). SVs with allelic frequencies > 0.7 in the germline, which we considered clonal, tended to be detected at stable levels in primary and recurrent tumors (**Fig. S4E-H**). The subclonal nature of some SVs in the germline, and their fluctuations in primary and recurrent tumors, were unexpected findings and we set out to further substantiate these observations.

In order to explore the observed SVs with an independent computational method, we re-called SVs with the Genome-wide Reconstruction of Complex Structural Variants (GROC-SVs) package *(34)*. We subset SV predictions to those for which we had sufficient physical coverage in all samples (see Methods), thus retaining SVs supported by at least 200 barcodes (i.e. DNA molecules) spanning each breakpoint (**Supplemental file 3**). GROC-SVs models noise specific to 10X linked-read data and assigns both a binary state corresponding to the presence or absence of a breakpoint in each sample, as well as a p-value to the likelihood of detecting that breakpoint given the local read support. We stratified SVs in each patient based on their pattern of presence or absence across samples. For instance, somatic SVs were absent from the germline and appeared in one or more tumor samples (**Fig. 4**). Germline events were detected in all samples, and in patients 1 and 3 could be further stratified into those inherited from a parent or occurring *de novo*. In all pGBM patients, we identified changes in SV allelic frequencies between tumor resections. Allelic frequency changes could represent loss or acquisition of somatic SVs in the tumors (**Fig. 4H**), and are often observed in our cohort to occur in genes with previous evidence for structural variation in pediatric cancers (**Fig. 4B,C,G,I**) *(35)*. For patient 1, we identified several *de novo* germline SVs (which were not detected in the parents) with allelic frequencies in the tumor samples (**Fig. 4C**). These data point to complex patterns of propagation of SVs from the germline to tumor samples, and highlight for the first time the presence of putative *de novo* germline SVs in pGBM patients.

**Fig. 4.**
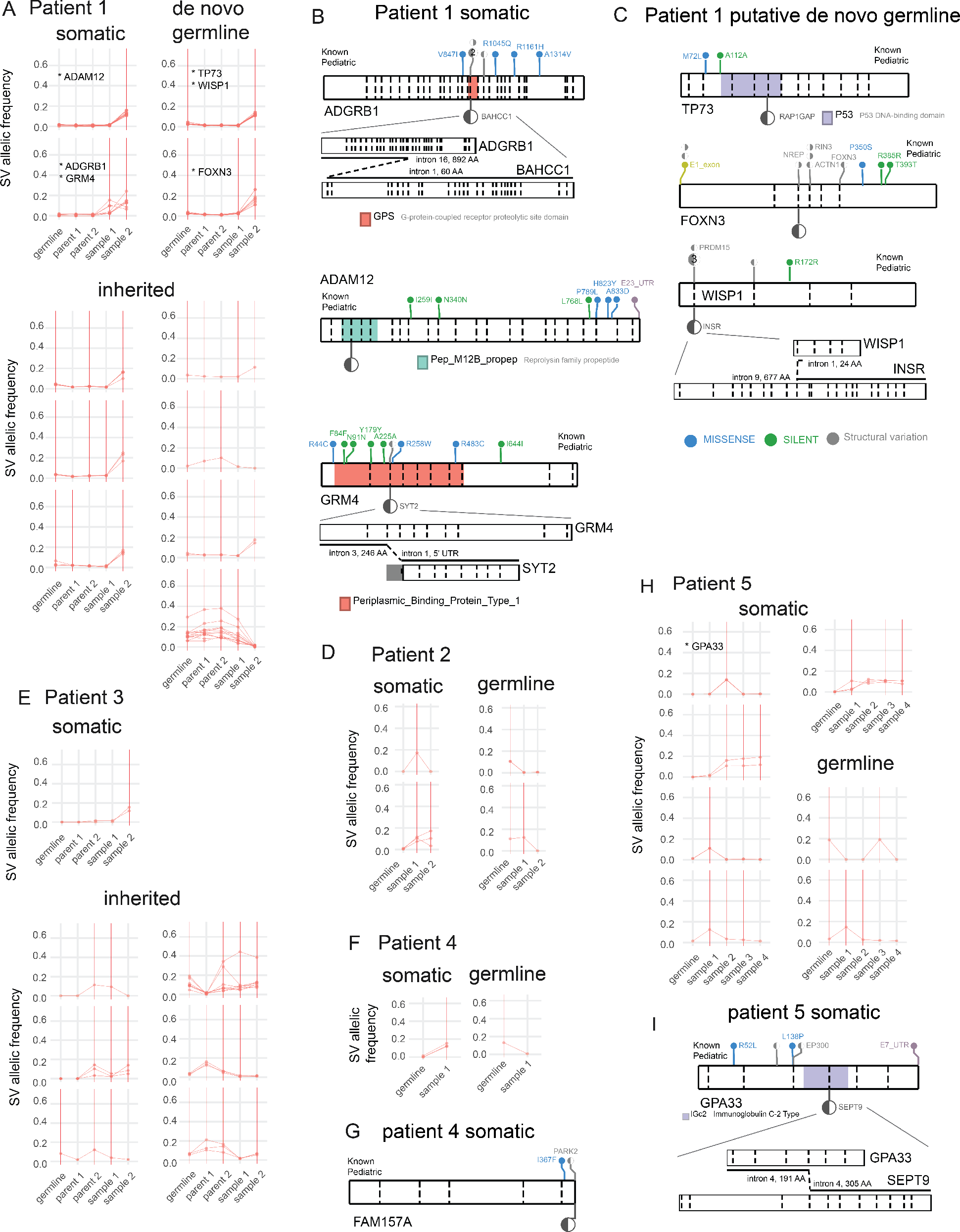
Dynamic contributions of structural variants to tumor genomes. Allelic frequencies for structural variants called by GROC-SVs are plotted across samples for all patients (patient 1 (A), patient 2 (D), patient 3 (E), patient 4 (F), and patient 5 (H)). Red lines connect dots that represent the allelic frequency of 10x barcodes supporting a particular SV, such that higher values represent a higher level of support for the SV. Only SVs with a minimum of 200 barcodes in each patient sample are shown. Red vertical lines highlight samples for which an SV was confidently called in the corresponding sample. Inset gene names represent genes shown in panels B, C, G, and I. Samples include the germline for primary tumors and their recurrences, and in germlines of the parents of patient 1 and patient 3. In these patients, germline events are stratified into inherited vs *de novo* events, and those observed in both parental germlines were excluded unless they were absent in another sample. Panels (B), (C), (G), and (I) contain gene level diagrams of select SVs that occur in genes previously associated with pediatric cancers. Gene diagrams include exon boundaries (dashed horizontal lines), protein domains (coloured boxes, only including domains overlapping the SV breakpoint), previously annotated somatic alterations in pediatric cancers (lollipops above the gene diagram), and the breakpoint observed in a particular patient (larger lollipop below the gene diagram). For SV events spanning genes ADGRB1-BAHCC1, GRM4-SYT2, and GPA33-SEPT9, the predicted effect of the structural variant on the structure of the genes is shown below the main gene model.

### Subclonal deletions at the*ATRX* locus in germlines and pGBM

The data organization principles underlying linked-read WGS data facilitated identification and visualization of subclonal SVs. Among these SVs, we identified large deletions in *ATRX* that removed almost the entire coding region of the gene (**Fig. 5A**). Somatic SNVs in *ATRX* are known to occur in pGBM *(3, 36)*, and loss of ATRX protein is routinely tested in GBM specimens in the clinic with immunostaining techniques. Our data indicate that *ATRX* harbors both SNVs and SVs in pGBM. Just outside the deletion breakpoints, there are a series of Alu and SINE repetitive elements, and recombination between these repetitive sites may be a mechanism behind the generation of the *ATRX* deletion. Of note, we detected putative subclonal *ATRX* deletions in the germline of pGBM patients. In order to validate the presence of putative *ATRX* deletions with an independent approach, we designed a TaqMan probe for copy number assays that targets the putatively deleted region (“ATRXdel” probe; **Fig. 5B**). A control TaqMan probe was designed to target a region on chromosome 5 predicted to be diploid based on our WGS data (“control” probe; see Supplemental Material). We then performed copy number TaqMan assays using genomes from both the germline and the tumors of our pGBM patients. TaqMan signal for the ATRXdel probe was internally normalized to signal for the control probe and number of X chromosomes. A second normalization was performed against the control diploid genome. This TaqMan assay confirmed the *ATRX* deletion in both germlines and tumors of all pGBM patients with the exception of patient 5, where the deletion was only detected in sample 2 (**Fig. 5C** and **Fig. S5**). In order to validate these findings in an independent cohort, we performed WGS with linked reads on three more pGBM samples and their matched germlines collected in Toronto. Our data show evidence of subclonal deletions at the *ATRX* locus in all samples (**Fig. S6A**). These findings confirm our WGS and copy number data and show that subclonal deletions in *ATRX* exist in both the germline and tumors of most pGBM patients.

**Fig. 5.**
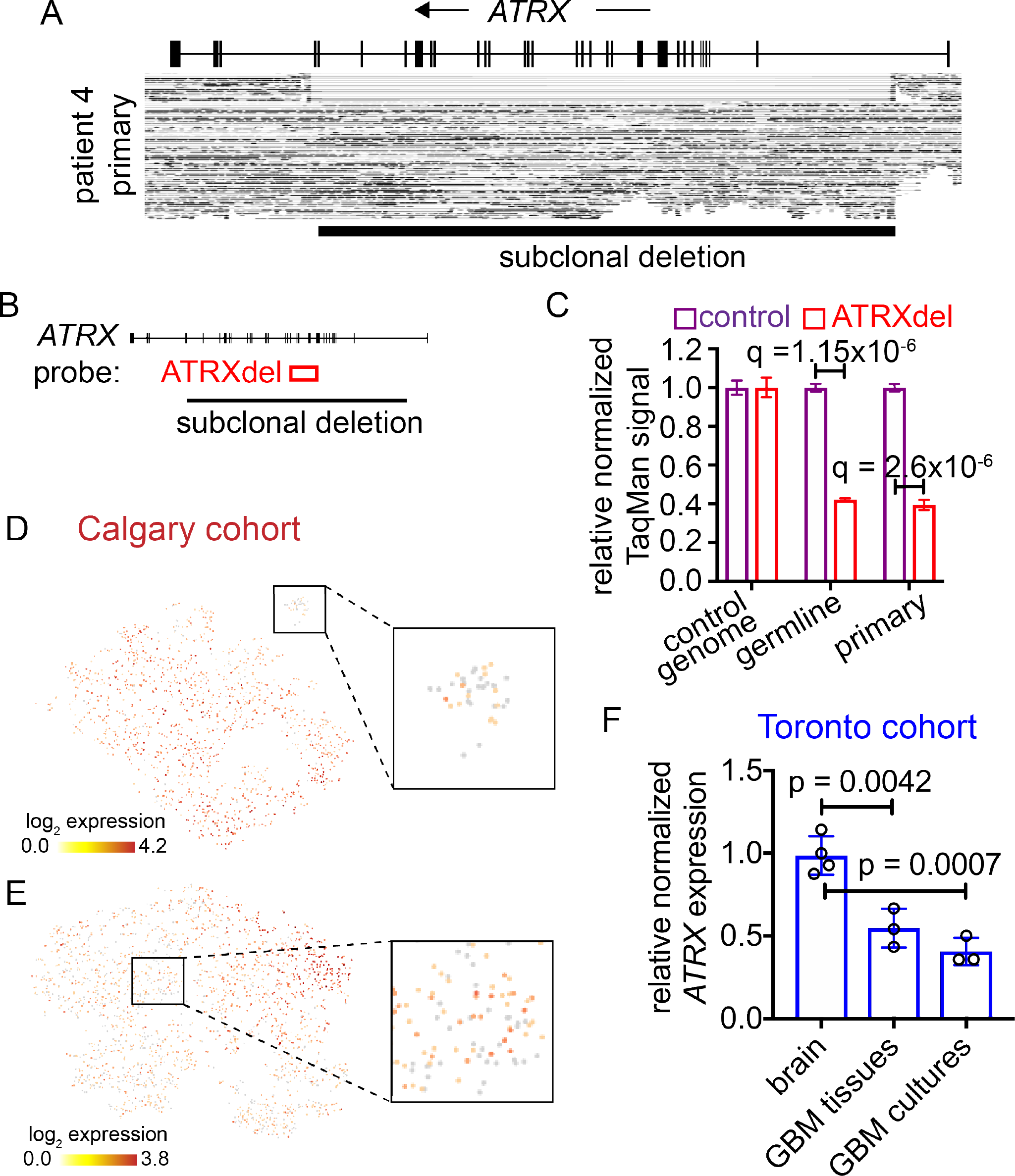
Subclonal *ATRX* deletions in the germline and tumors of pediatric GBM patients. (A) Putative subclonal deletions at the *ATRX* locus in tumors from a pediatric GBM patient. (B) Diagram describing the binding locations of a TaqMan probes (ATRX del) designed to detect copy number variations at the *ATRX* locus. (C) TaqMan assays to detect copy number variation at the *ATRX* locus on samples for patient 4. (D) Single-cell RNA-seq in a xenograft derived from patient’s 3 recurrence. *ATRX* expression levels are represented in this tSNE plot. (E) Single-cell RNA-seq data for recurrence 3 of patient 5. This tSNE plot displays expression levels for *ATRX*. (F) RNA-seq data for *ATRX* expression in non-neoplastic brain tissues (n = 4), bulk pediatric GBM samples (n = 3) and their matched primary cultures (n = 3). *ATRX* expression was normalized to the average for non-neoplastic brain tissues.

We expected that subclonal *ATRX* deletions may result in heterogeneous expression of the gene in populations of tumor cells. Since our longitudinal collection was assembled over a long period of time, storage of pGBM samples was not compatible with single-cell RNA-seq (scRNA-seq) methods. We therefore decided to assess transcriptional heterogeneity using patient-derived xenografts (PDXs), which were available for sample 2 of patient 3 and for sample 4 of patient 5. We successfully generated scRNA-seq data for 1,327 cells from patient 3’s sample 2, and for 2,204 cells patient 5’s sample 4. As predicted from the WGS data and TaqMan assays, scRNA-seq data showed heterogeneous expression of *ATRX* in both PDXs (**Fig. 5D,E**). In contrast to other genes, for instance *H3F3A* and *H3F3B* (both of which encode the histone variant H3.3), were more homogeneously expressed (**Fig. S6B,C**). Widespread decreased *ATRX* expression in pGBM compared to non-neoplastic brain tissue (n = 4) was confirmed by RNA-seq of bulk pGBM tissue (n = 3) and their matched primary cultures from an independent cohort collected in Toronto (**Fig. 6F**). Furthermore, analysis of an independent genomic dataset confirmed that subclonal *ATRX* mutations are detected in the tumors of pGBM patients (**Fig. S6D**). Our data therefore show that in addition to somatic SNVs in *ATRX*, pGBM patients may also carry subclonal *ATRX* deletions in both their germlines and tumors.

**Fig. 6.**
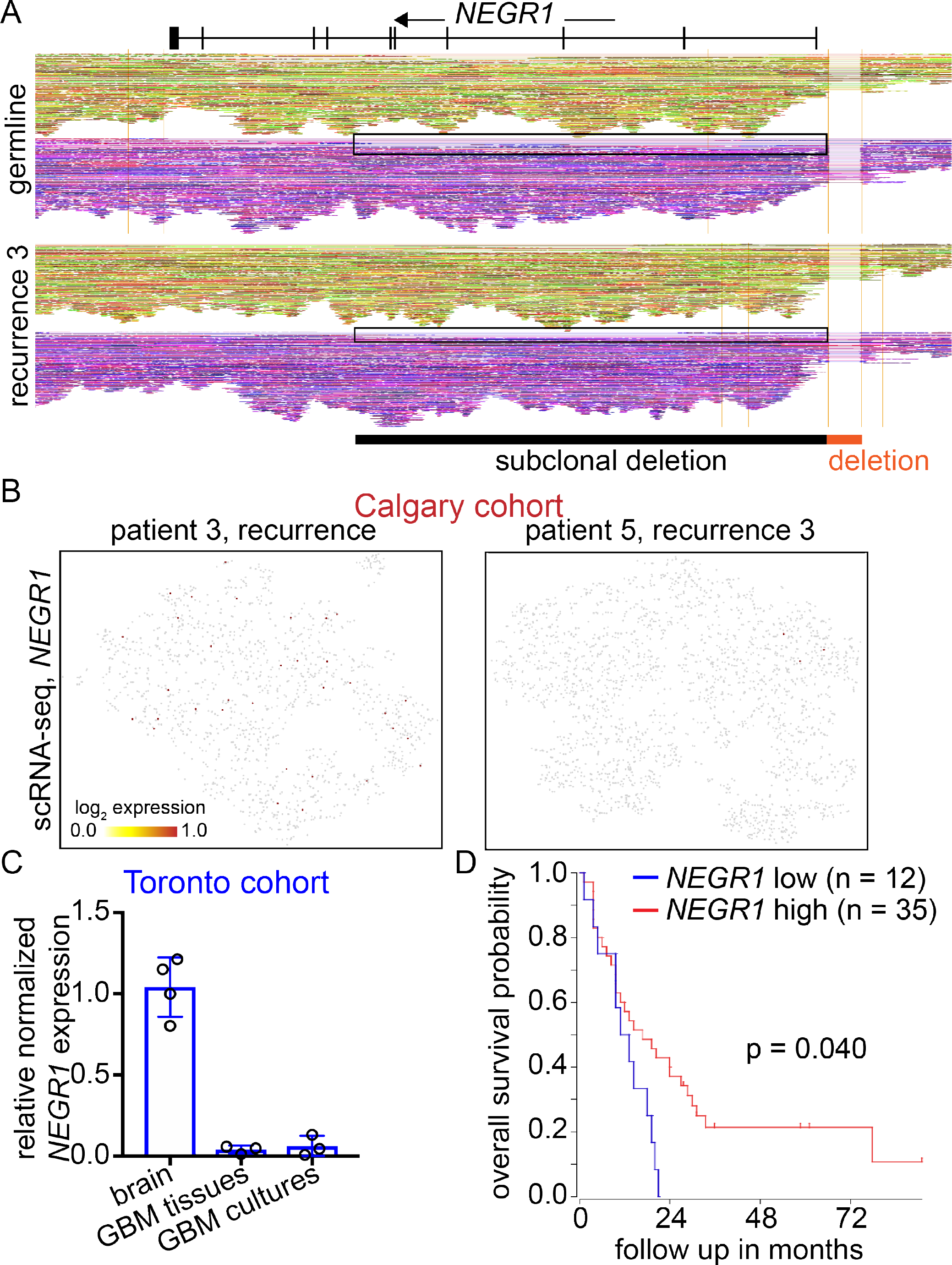
Germline deletions at the *NEGR1* locus are common to all patients. (A) Coverage of linked-reads at the *NEGR1* locus. Linked-reads are grouped based on haplotypes (red/green hues for one haplotype, purple/violet for the second). A deep deletion is observed in the region immediately upstream of the gene for the germline and the third recurrence for a pediatric GBM patient. A subclonal deletion that extends into the gene body is also visible (black box). (B) Single-cell RNA seq for a pGBM xenograft derived from the recurrence of patient 3, and a xenograft derived from the third recurrence of patient 5. Expression levels of *NEGR1* are shown. (C) RNA-seq data for *NEGR1* expression in non-neoplastic brain tissue (n = 4), bulk GBM tissues (n = 3) and their matched GBM primary cultures (n = 3). (D) Kaplan Meier curve for pediatric GBM patients in the Paugh dataset, stratified by expression of *NEGR1*. Log-rank p = 0.040.

### Identification of conserved intergenic SVs in pGBM samples and germlines

Genome-wide coverage is a strength of our data compared to previously published datasets. We decided to leverage this asset by looking for intergenic SVs that were conserved in our pGBM samples. We found only one intergenic SV that was shared among all pGBM samples and their germlines, a deletion immediately upstream of *NEGR1* (neural growth regulator 1; **Fig. 6A**). This upstream deletion was heterozygous or homozygous in different patients, but the breakpoints were absolutely conserved. Some haplotypes displayed evidence of subclonal deletions in part of the gene (black box in **Fig. 6A**). Analysis of the GTEx database *(37)* showed that *NEGR1* is expressed at higher levels in brain tissues - including frontal cortex and the cerebellar hemisphere - than in other anatomical locations (**Fig. S6E**). *NEGR1* mutations have been linked to obesity *(38)* and dyslexia *(39)*. In mice, *Negr1* enhances neurite outgrowth in cortical neurons, and *Negr1* knockout animals have axonal defects and behavioral deficits *(40)*. Expression of this gene is downregulated in different cancer types, and downregulation of *NEGR1* promotes migration *(41)*. Our scRNA-seq show that PDX samples with deletions upstream of *NEGR1* have undetectable expression of this gene in nearly all cells (**Fig. 6B**), suggesting that the SV affects regulatory regions. We observed strong repression of *NEGR1* in an independent set of pGBM samples and their matched pGBM primary cultures from Toronto (**Fig. 6C**). Importantly, our analysis of a previously published pGBM cohort *(42)* shows that *NEGR1* expression can stratify pediatric glioma patients based on overall survival, with tumors expressing lower levels of the gene having worse prognosis (log-rank p = 0.040; **Fig. 6D**).

### Evidence for the existence of relatively quiescent cancer stem cells in pGBM

We questioned whether the observed variability of the mutational profile might correlate with proliferative heterogeneity in the respective contributions made by subclones to tumor growth and recurrence. Recently, cell tracing studies based on the genetic barcoding of freshly-isolated adult GBM cells transplanted into PDX mouse models have suggested that tumor cells may adhere to a common fate behavior, reminiscent of a normal neurogenic program *(43)*. Specifically, a signature negative binomial dependence of the distribution of variant allele frequencies (VAFs) provides evidence of a conserved proliferative hierarchy in which slow-cycling tumor stem cell-like cells give rise to a rapidly cycling, self-renewing, progenitor-like population (**Fig. 7A**) (Extended Data Figure 4 of *(43)*). Applied to the patient samples in the current study, we find that the VAF distribution obtained from tumor samples is also largely consistent with a negative binomial dependence (**Fig. 7B-E** and **Fig. S7A-I**), suggesting pGBM tumor cells may be defined by a similar proliferative hierarchy (see Methods). This conclusion is reinforced by focusing on the ensemble of *de novo* point mutations that are acquired, or only rise above the detection threshold, during recurrence (**Fig. S7J-N** and Methods). By contrast, mutations that are shared between the primary and recurrent tumor samples show VAFs that are peaked at larger values, consistent with the predominance or fixation of subclones within the tumor population (**Fig. S7O-R**). Our data from our pGBM patient cohort are therefore consistent with a hierarchical model for pGBM, characterized by slow-cycling cancer stem cells that give rise to fast-proliferating progenitor-like cells, which eventually produce non-dividing “differentiated” cells.

**Fig. 7.**
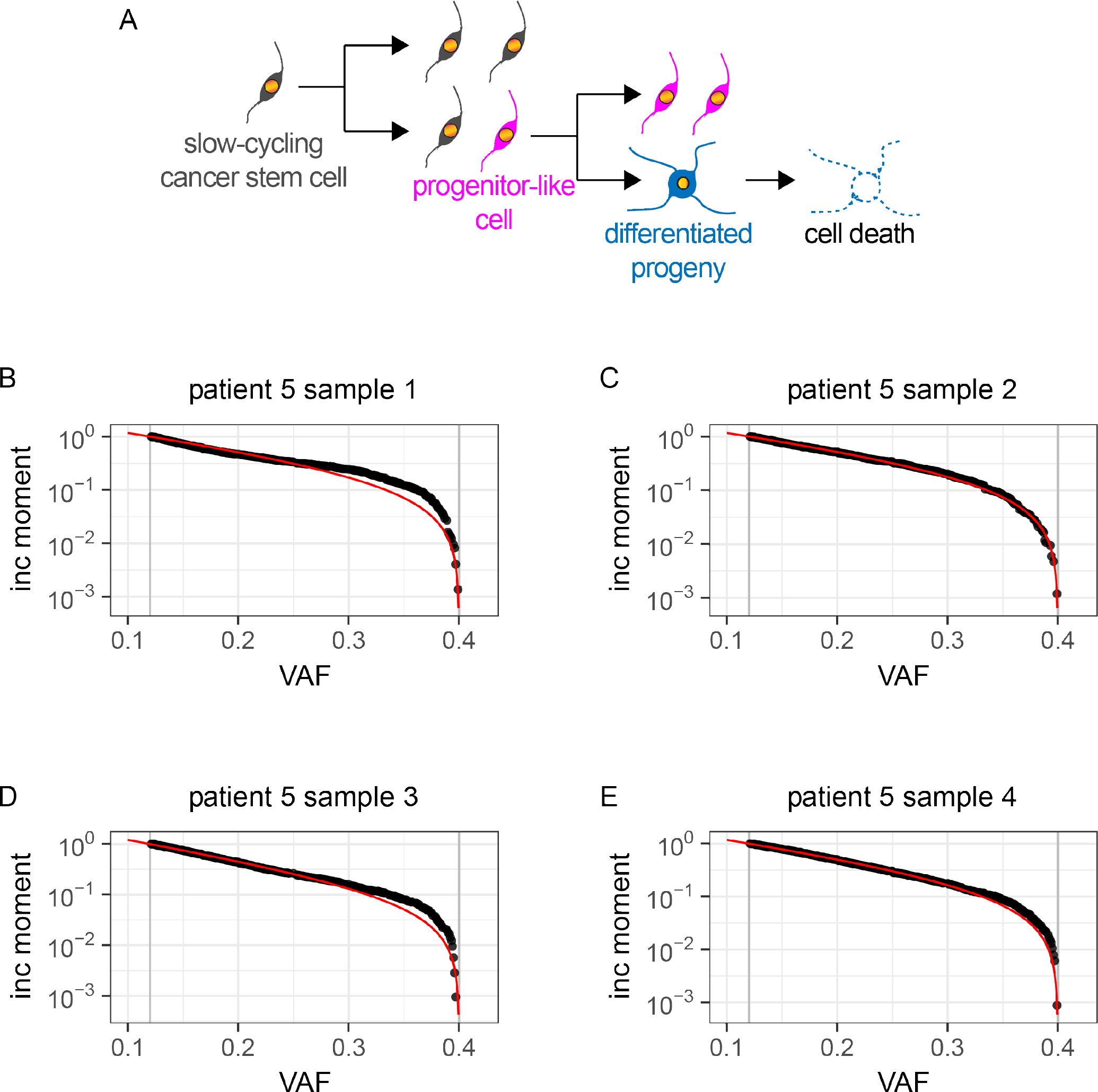
Evidence for a hierarchical organization of pGBM. (A) Diagram illustrating a testable model for pGBM organization and growth. Under a hierarchical model that includes relatively quiescent cancer stem cells giving rise to proliferating progenitor cells and non-proliferating cells, the incomplete moment of variant allele fractions (VAFs) would follow a negative binomial distribution. (B-E) Fit of the first incomplete moment of the VAF distribution based on a neutral hierarchical model of tumor cell dynamics (red line) to data obtained from patient 5 (black points). The grey lines indicate allele frequency cut-offs due to sequencing resolution (lower) and ploidy (upper).

## Discussion

Cancer is often compared to a caricature of developmental processes. Molecular profiling of pGBM suggests that this cancer has features that make it appear like a caricature of hereditary disorders. Notwithstanding large differences in terms of mutated genes among tumors, in every patient we identified somatic SNVs in several genes linked to hereditary conditions, including MeCP2, which is linked to Rett syndrome *(44)*, and autosomal dominant genes linked to severe intellectual (*ZMYND11*, OMIM number 608668) and developmental syndromes (*DSE*, linked to Ehlers-Danlos syndrome; *REEP1*, OMIM number 609139). Anecdotally, we found somatic SNVs in numerous genes associated with blood disorders, such as the recurrent SNVs in *VWF* (a gene linked to von Willebrand disease), *LMO2* (leukemia), *CR2* (systemic lupus erythematosus), *ASXL1* (myelodysplastic syndrome), *ITAGA2B* (bleeding disorder), and *MLL1* (leukemia). Genetic lesions did not cluster in specific pathways or gene classes, but this might reflect our small sample size. Interestingly, the pediatric brain tumor diffuse intrinsic pontine glioma has recurrent somatic mutations in *ACVR1 (24, 45–47)*, and germline mutations in this gene lead to the hereditary syndrome fibrodysplasia ossificans progressiva *(48)*. The detection of germline variants in pGBM patients, and the identification of somatic SNVs in genes associated with hereditary syndromes, give pGBM a characteristically different molecular flavor from adult GBM. Because of the early onset of pGBM, studies of larger cohorts will be required to determine if germline and somatic events occurring during embryonic and/or fetal development cooperatively contribute to pGBM initiation and progression. These studies will be particularly important given that currently no genetic or environmental factor has been associated with sporadic pGBM. It would be important to determine whether the *de novo* germline variants identified in this study arise in the germ cells of the parents during the normal mutagenic processes of genome replication *(49)*, or if they arise during early embryonal or fetal development. Our data showing that some variants have low (< 0.2) allelic frequency in the germline of pGBM patients suggest that these individuals might display somatic mosaicism resulting from mutational processes during postzygotic stages of development. Interestingly, clonal hematopoiesis has recently been shown to be associated with a 10-fold increase in risk of developing hematologic cancer *(50, 51)*, and was observed in both adult and pediatric patients with non-hematologic cancers *(52)*. In pediatric cancer patients, clonal hematopoiesis was identified in 5-10% of individuals, with highest frequency in children aged 0-9 years. There is therefore a possibility that some subclonal SVs with < 0.2 allelic frequencies in the germline that became barely detectable or undetectable in tumors might reflect clonal hematopoiesis. In this scenario, the frequency of germline SVs in the tumor may depend on the extent of tumor infiltration of hematopoietic cells carrying that specific SV. However, we identified this phenomenon in all pGBM samples, which is more than expected based on available published data *(52)*. The low AF of some germline SVs is therefore more likely the result of developmental processes that selected them over time.

By sequencing the germlines of trios, our data provide an important perspective on the developmental origins of pGBM. Our data show that some potentially deleterious germline variants (like a *TP53* exonic deletion in patient 1) can be inherited from a healthy parent. At the same time, we observed *de novo* non-synonymous SNVs in the germlines of both patient 1 and patient 3, as well as *de novo* germline SVs in patient 1. The expected rates of emergence of *de novo* non-synonymous SNVs is very low, based on recent literature evidence on trios that included probands affected by disorders with high heritability. Trio studies in the context of sporadic autistic spectrum disorder identified 181 *de novo* non-synonymous SNVs in 189 probands, which corresponds to an average of 0.96 *de novo* non-synonymous SNV per child *(53)*. A second trio study found 32 *de novo* non-synonymous SNVs in 53 probands with schizophrenia, corresponding to a rate of 0.60 *de novo* non-synonymous SNVs per individual *(54)*. In the same study, 7 out of 22 unaffected individuals (control group) had *de novo* SNVs, with 4 out of 7 SNVs being non-synonymous. While acknowledging that our sample set is small and that there may be differences in numbers of SNVs detected and called based on sequencing platform and computational pipelines used, our data offer proof-of-principle that mutational processes are at play in the parents’ germ cells or in the early developing zygote leading to *de novo* germline variants in at least some pGBM patients. Because of the rare nature of pGBM, and difficulties in re-contacting families of pGBM patients, the trios we generated represent extremely valuable resources. In the future, it will be important to assemble germline WGS data on trios of pGBM probands in a systematic fashion, in order to assess and investigate genomic predisposing events.

Using WGS data with linked reads, we have identified a set of germline SVs that tend to occur at high frequency in our cohort. One of these examples is subclonal deletions at the *ATRX* locus in both germlines of four out of five patients and tumor tissues of all patients. Recent work showed that deletion of *Atrx* in mouse neural progenitor cells or *ATRX* in adult glioma stem cells alters patterns of chromatin openness genome-wide and leads to enhanced cell migration *(55)*. Because of the established involvement of *ATRX* in gliomagenesis, we speculate that *ATRX* deletions might be a predisposing genetic lesion for pGBM. Germline deletions upstream of *NEGR1* were present in every pGBM patient, in both the Calgary and in the Toronto cohorts. These deletions had conserved breakpoints, and were either homozygous or heterozygous in different patients and resulted in loss of *NEGR1* expression. Given that this gene has an important role in neuronal development in mouse models *(40)*, and its decreased expression positively affects migration *(41)*, it is tempting to speculate that loss of *NEGR1* expression may confer a competitive advantage at early stages of tumor development. Lesions in *ATRX* and *NEGR1* may cooperate with other SVs we identified in the germlines of pGBM patients to determine a certain level of predisposition to the development of cancer. For instance, we found that 4 out of 5 patients also harbored germline deletions at the *BTNL8/3* locus (**Fig. S3A**), causing loss of expression of the affected genes (**Fig. S3B,C**). *BTNL8* encodes a protein that activates immune cells *(32, 56)*. We hypothesize that multiple lesions in genes involved in brain development and function (eg *ATRX* and *NEGR1*) and in activation of the immune system (eg *BTNL8*) might provide a selective advantage to cells that participate in the early steps of the tumorigenic process.

Importantly, our analyses show that pGBM is characterized by intratumoral genetic heterogeneity and subclonal architecture, which rapidly evolves at recurrence. Our work, together with previous reports on pediatric medulloblastoma *(15)* and diffuse intrinsic pontine glioma *(11)*, indicates that pediatric brain cancers have more complex genetic architectures than previously thought. The genetic divergence between diagnostic and recurrent samples should be kept into consideration when designing personalized treatment approaches. In addition to genetic heterogeneity, our data provide evidence for the existence of a functional hierarchy in pGBM. VAF distributions in pGBM samples are consistent with a hierarchy comprising relatively quiescent cancer stem cells, fast-proliferating progenitor cells and non-self-renewing and non-dividing cells destined to undergo cell death. Single-cell barcoding experiments with xenografts derived from adult GBM patients supported a similar hierarchy *(43)*. Although adult GBM and pGBM are molecularly different, hierarchical organization may be a common feature of both. If relatively quiescent, self-renewing cancer stem cells reside at the apex of a pGBM hierarchy, these cells might be less sensitive to radiation and antimitotics used to treat this cancer, explaining why recurrences did not have a significantly higher mutational burden than the diagnostic samples.

Overall, our data provide strong evidence for the existence of subclonal architecture in pGBM and relatively rapid evolution at recurrence, irrespective of treatment. Reconstruction of mother-father-patient trios revealed the potential for inherited and *de novo* germline variants and somatic mutations to contribute to the etiology of this pediatric cancer.

## Methods

### Contact for reagent and resource

Further information and requests for resources and reagents should be directed to and will be fulfilled by the Lead Contact, Marco Gallo. Sharing of primary samples, based on availability, may be subject to MTAs and will require appropriate research ethics board certifications.

### Human samples

All samples were collected and used for research with appropriate informed consent and with approval by the Health Research Ethics Board of Alberta and the Research Ethics Board of the Hospital for Sick Children. Fresh tumor tissue and blood samples from each patient were collected by the Clark Smith Brain Tumour biobank at the University of Calgary and the Hospital for Sick Children and preserved at either −80°C or in vapor phase liquid nitrogen. For selected cases, a portion the fresh tumou material was also allocated for patient-derived xenograft preparation (see below). Anagraphical information of sample donors are in **Table S2**and **Table S3**.

### Genomic DNA extraction and library preparation

High molecular weight genomic DNA (gDNA) was extracted from frozen tumor tissue and frozen blood samples with the Qiagen MagAttract^®^ HMW DNA Kit (catalog # 67563). gDNA fragment size and distribution were quantified with the Agilent Technologies 2200 TapeStation Genomic DNA Assay. Samples with gDNA fragment size > 50 kb were used for library construction. The 10xGenomics Chromium™ Genome Library Kit & Gel Bead Kit v2, 16 reactions (catalog # PN-120258) was used for library preparation. Size and distribution of all sequencing libraries were quantified with the Agilent Technologies 2200 TapeStation D1000 Assay.

### Whole-genome sequencing and linked-read data analysis

Whole-genome sequencing was performed at The Centre for Applied Genomics at the Hospital for Sick Children (Toronto, ON) with a HiSeq X Series (Illumina) instrument. Libraries were validated on a Bioanalyzer DNA High Sensitivity chip to check for size and absence of primer dimers, and quantified by qPCR using Kapa Library Quantification Illumina/ABI Prism Kit protocol (KAPA Biosystems). Validated libraries were paired-end sequenced on an Illumina HiSeq X platform following Illumina’s recommended protocol to generate paired-end reads of 150-bases in length. Each library was loaded on a single lane. All libraries were sequenced at 30x coverage. Post-sequencing, fastq files were analyzed with the package LongRanger-2.1.6 (10xGenomics) on servers hosted by the Centre for Health Genomics and Informatics (University of Calgary). LongRanger performed alignment of reads to the reference genome (GRCh38), assembled linked-reads, generated haplotype blocks, and called single nucleotide polymorphisms and structural variants. LongRanger output files were visualized with Loupe browser 2.1.2 (10xGenomics). SNVs were called using both Mutect2 *(57)* and Strelka2 (https://github.com/Illumina/strelka). Only SNVs called by both methods were included for downstream analysis. Annotation of SNVs was performed using ANNOVAR *(58)*.

### Structural variant calling using GROC-SVs

GROC-SVs was run for all samples in each patient independently using default parameters. SVs in patients with parental germlines available were further analyzed in order to empirically infer a cutoff for barcode coverage that was sufficient to call an SV. For instance, events observed in both parental germlines and the patient tumor samples but not in the patient germline must be false negative calls in the patient germline as they are inherited. A threshold of 200 barcodes (physical coverage) was sufficient to minimize false negative calls across the cohort, without significantly compromising our ability to detect all true positive events. For patients 1 and 3, events observed in both parents were excluded from further analysis, unless they were somatically altered (i.e. absent in the primary or recurrent tumor). The overlap of SV breakpoints and genes was assessed using bedtools intersect, and these breakpoints were visualized using the ProteinPaint software (pecan.stjude.cloud/proteinpaint) along with known pediatric aberrations from a number of databases. These include PCGP (St. Jude - WashU Pediatric Cancer Genome Project), TARGET (Therapeutically Applicable Research To Generate Effective Treatments), SCMC (Shanghai Children’s Medical Center pediatric ALL project), UTSMC (UT Southwestern Medical Center Wilms’ tumor study), and DKFZ (German Cancer Research Center Wilms’ tumor study) *(35)*.

### Somatic SNV detection and filtering using varscan2

Somatic variants were also called genome-wide using varscan2 (v2.4.3). Varscan2 tumor-normal variant calling produced variants categorized as somatic, germline or loss of heterozygosity (LOH). Briefly, alignments from 10X Genomics LongRanger underwent tumor-normal variant calling using samtools mpileup (v1.7), to produce a single tumor-normal mpileup file *(59)*. The mpileup files were processed to yield variants using the varscan2 <u>somatic</u> function and high confidence variants were isolated using the varscan2 <u>processSomatic</u> function. Bam read counts files were created using a combination of tumor or normal bam and the appropriate high confidence variant coordinates; one for LOH, one for germline and one for somatic variants. Variants were filtered using the <u>fpfilter</u> function under dream-3 settings to produce a high confidence and filtered set of variants.

### Somatic copy number alteration (CNA) calling, filtration and segmentation

Somatic copy number alterations were estimated using varscan2. Tumor-normal mpileup files (see SNV detection, above) were used to generate raw copy numbers using the <u>copynumber</u> function, and further adjusted for for GC content using the <u>copycaller</u> function. Adjusted calls underwent segmentation using a recursive algorithm, circular binary segmentation as implemented in the package DNAcopy *(60)*. Segmented CNA was analyzed and readjusted using <u>copycaller</u> with <u>recenter-up</u> or <u>recenter-down</u>, for segments showing a consistent deviation from a baseline (0.0). Adjacent segments of similar copy number log ratios (<0.25 difference) were merged using a varscan2 affiliated perl script and a custom bash script.

### Somatic SNV classification using mclust

Variants were classified based on their variant allele frequencies (VAFs) into distinct clusters using the R package mclust *(61)*. Mclust uses machine-learning model-based estimation based on finite normal mixture modelling using Expectation-Maximization (EM) and the Bayesian Information Criterion, essentially allowing for the grouping of variants based on allele frequency into distinct distributions. These can be manually assigned into “homozygous’, “clonal’ or “subclonal’.

### Cancer Cell fractions (CCF)

Variant cancer cell fractions (CCF) indicates the prevalence of a mutation throughout the tumor sample, with higher CCF values indicating variants clonal in nature. CCFs were calculated as follows *(62)*:

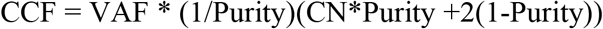

Where VAF is the variant allele frequency, CN is the local copy number of the mutation and purity is the estimation as calculated by EXPANDS. Both VAF and CN were calculated using varscan2 (see above).

### Phylogenetic reconstruction from combined SNV and CNA data

CNA and loss-of-heterozygosity (LOH) data were combined with somatic variants to infer subclonal architecture using EXPANDS *(22)* independently for each sample. EXPANDS (V2.1.1) was run using the runExpands function with default settings. As per the default, only subpopulations that had a minimum cellular frequency (CF) of 0.1 or greater were taken into consideration. Mutations that were not clusterable were excluded from reconstruction. Mutations were assigned to all appropriate populations and subpopulations, (ie a mutation can be assigned to a parent population (trunk) and a “daughter’ subpopulation (branch).

Phylogenetic reconstruction of the subpopulations in primary and recurrent tumors was performed by calculating the Manhattan distance metric between all subpopulations, followed by hierarchical clustering using full linkage. Visualization of the phylogenies were created using the as.phylo function from the ape package *(63)*.

### Copy number spectrum plots

CNA data from varscan2 copy number analysis was used to visualise copy number gains and losses. The copy number spectrum function for the GenVisR package provides a method of visualizing primary and recurrence CNA changes *(64)*. The genome boundaries were created using BSgenome.Hsapiens.UCSC.hg38 reference.

### Loss of heterozygosity plots

Germline and loss of heterozygosity (LOH) variants called by varscan2 were used to visualize LOH regions of the genome using the lohspec function in the GenVisR package. This enabled detection of copy-neutral LOH regions where one parental locus is deleted and the other is duplicated. LOH regions were found using a 2.5 Mbp sliding window to calculate difference in variant allele frequency of germline and LOH heterozygous variants in tumor and normal samples. Within each window the absolute VAF difference is calculated and a mean is obtained. The algorithm continues recursively by moving the windows forward 1 Mbp and calculating the mean tumor-normal VAF difference of LOH and germline variants genome-wide, until a stable solution is found.

### Mutational signatures

To determine if different mutational processes were occurring we analyzed mutational signatures. Sets of somatic variants from distinct subclonal populations in primary or recurrent tumors identified by EXPANDS were analyzed separately, and the mutation signatures were determined using the deconstructSigs (v1.8.0) package *(65)*. Mutational signatures were generated for each subclonal population group, all groups were above 50 variants, with the exception of 3f (patient 3), 1ef (patient 1) and 2b (patient 2). DeconstructSigs was run using 30 signatures from the Catalogue Of Somatic Mutation In Cancer public repository as the reference signatures *(66)*. Default settings were used when generating mutational signatures with the exception of genome normalization and weight signature cutoff, which was changed to 0.05.

### Allele frequency distribution

Previously, it has been argued that the dynamics of tumor cells may find a signature in hallmark statistical clone size dependences *(67, 68)*. Specifically, for systems characterized by a proliferative hierarchy in which slow-cycling tumor stem-like cells give rise to rapidly cycling, self-renewing, progenitor-like cells that in turn generate short-lived non-cycling cells, the distribution of variant allele fractions (VAFs) is predicted to acquire a negative binomial frequency dependence *(43, 67)*:

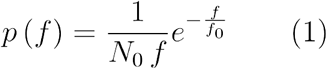

Here, the leading *1/f* dependence, highlighted by Williams et al. 2016, represents a generic signature of neutral clone competition. To explore whether the current tumor WGS data can be embraced within the same paradigm, we considered the VAF distribution from patient samples. To this end, we considered a fit to the first incomplete moment of the VAF distribution, which for a negative binomial size dependence takes a pure exponential form *(43, 67)*.

Due to the sequencing resolution limit (~150x physical coverage) we introduced a lower cut-off on the allele frequencies at *f*_*min*_ = 0.12 *(69)*. At the upper end of the scale, we set a cut-off at *f*_*max*_ = 0.4, noting that, in the absence of copy number variation, mutations with an allele fraction of 0.5 and above will include those that are fixed across the population. We therefore adapted the expression for the first incomplete moment to account for this limited allele frequency interval of [*f*_*min*_, *f*_*max*_], defining the measure,

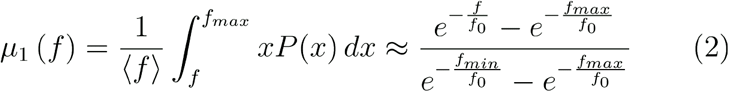

where < *f* > denotes the average allele frequency within the interval and *f*_0_ is the fitting parameter. For details on how the VAF distribution is derived, we refer to ref. *(67)*.

The *NonlinearModelFit* function in *Mathematica 11.0.1.0* was then used to perform a least-squares fit of the first incomplete moment to the sample (allele frequencies corrected for the modest degree of copy number variation and purity). The results of the fit can be found in Table S7, Figure 7B-E and Figure S7A-D. Overall, the findings are largely consistent with a negative binomial tumor clone size dependence. These results suggest that tumor cells are organized in a conserved proliferative hierarchy, with tumor growth characterized by neutral competition between constituent tumor subclones.

For completeness, we then considered a direct fit of the VAF distribution to a negative binomial size dependence (1) over the same interval. Grouping the data into bins of size (*f*_*max*_ − *f*_*min*_)/30, the results of a least-squares fit, obtained using the *NonlinearModelFit* function in *Mathematica 11.0.1.0*, is shown in **Fig. S5E-I**.

### Evolution of allele fractions across recurrences

To further challenge the hierarchical model tumor cell dynamics, we took advantage of the availability of recurrent tumor samples. Within the framework of the model, competition between individual tumor stem cell clones is predicted to be neutral, allowing clones to readily adjust, up or down, their fractional tumor contribution. In this case, the ensemble of clones bearing mutations that are acquired, or rise above detection threshold, during recurrence provides a pure sample on which to test the model through the predicted negative binomial dependence of the VAF distribution. By contrast, mutations that are shared between primary and recurrent tumors are expected to depart from this dependence, either because they are clonally fixed within the tumor population (with VAFs around 0.5), or because they occupy a significant fraction of the tumor at the start of the recurrent period of tumor growth. Notably, for *de novo* mutations that arise at recurrence, using the same least-squares fitting procedure, we find a strikingly good agreement with a negative binomial dependence (**Fig. S5J-N**), while the distribution of shared mutations is typically peaked around larger values, as expected (**Fig. S5O-R**).

### Copy number assays for the*ATRX* locus

Custom TaqMan^®^ Copy Number Assay Probes (catalog # 4400294) from Applied Biosystems™ by Thermo Fisher Scientific were used to detect changes in copy number at the *ATRX* locus in all patients. Quantification was performed according to manufacturers’ protocols. Briefly, for each patient, a control DNA sample, patient germline DNA, primary tumor DNA, and recurrent tumor DNA were diluted to 5 ng/μL. DNA was quantified in triplicates with the Qubit dsDNA HS Assay Kit (catalog # Q32851). The TaqMan^®^ Genotyping Master Mix and TaqMan^®^ Copy Number Assay was added to each well of a 96-well plate. 20 ng of DNA was added in triplicate wells of a 96-well plate for each assay condition. The samples were then ran on a Bio-Rad CFX Connect™ Real-Time PCR Detection System with the following parameters: hold at 95°C for 10 mins, 40 cycles of 95°C for 15 sec and 606C for 60 sec. Fluorescence signal for the *ATRX* deleted region (ATRXdel probe) was first normalized to the fluorescence signal from the control diploid region probe. Fluorescence was additionally normalized to the diploid control DNA sample. TaqMan^®^ Probes are labeled with FAM at 5’ end and MGBNFQ at the 3’ end. For the probe in the *ATRX* deleted region; probe sequence: CACACCCAAATATTGGTAAAAAT, forward primer: TCCAGGACTTAGCAGGATGTGA, reverse primer: ACCCCATCAAGTAGATGGTAAGAAACT. For the probe in the control diploid region; probe sequence: ACACGGGTGTTGAAGACGC, forward primer: AGGCGCTGTGGAAACAATTATAGTA, reverse primer: TGTCCCCGTCAACGATCAC.

### Patient-derived xenografts

All mice were used with approval from the Animal Care Committee of the University of Calgary. Fresh tumor tissue was gently dissociated by trituration, filtered using a 0.7 μm filter, washed and resuspended in sterile DPBS (Gibco). Tumor cells were injected into the right striatum of the brains of CB17 SCID mice (Charles River Laboratory). Animals were monitored for tumor growth and tumors were isolated when signs of morbidity were observed. Initial tumor establishment for patient specimens SM4021 (recurrence of patient 3) and SM4058 (third recurrence of patient 5) took 59 and 41 days respectively. Tumors were then maintained by serial *in vivo* passaging in the brain of SCID mice every 1-2 months or cut into 1 mm x 1 mm tissue pieces and cryo-stored in 1 mL CryoStor CS10 solution (Stem Cell) for 10 minutes on ice prior to being stored at −80ºC. In experiments where cryo-stored tissue was used, tissue was thawed, dissociated into a single-cell suspension, filtered using a 0.7 µm filter and implanted into the brain of SCID mice. For single-cell sequencing, established xenografts were isolated and dissociated as described below. All xenografts were identity matched to the original patient material by short tandem repeat analysis using the AmpFLSTR Identifiler Plus kit (Applied Biosystems).

### Single-cell RNA-seq (Calgary cohort)

Xenograft tumor tissue was dissociated into single cells using the following protocol: 10 mL of ACCUTASE™ (catalog # 07920, Sigma-Aldrich) was added to tumor tissue for manual dissociation with surgical scissors. The sample was then incubated with sterile glass beads at 37°C, rotating continuously for 30 min. The glass beads were removed, and the sample was spun at 1000 rpm for 5 minutes at 25°C. The cell pellet was resuspended in neural stem cell media (NeuroCult™ NS-A Proliferation Kit (Human), catalog # 05751). Mouse cells were removed with the Miltenyi Biotec Mouse Cell Depletion Kit (catalog # 130-104-694) and the autoMACS^®^ Pro Separator and columns (catalog # 130-021-101). Protocol was followed as per manufacturers’ instructions. Viability and concentration of the purified human cells were quantified. Cell samples with viability > 80% were used for subsequent single-cell RNA-sequencing. ~2500 cells were loaded into the Chromium™ console and were used for library preparation. The 10xGenomics Chromium™ Single Cell 3’ Library & Gel Bead Kit v2, 16 reactions (catalog # PN-120237) was used for library preparation. Size and distribution of all sequencing libraries were quantified with the Agilent Technologies 2200 TapeStation High Sensitivity D1000 Assay. Single-cell RNA-sequencing was performed at the Centre for Health Genomics and Informatics at the University of Calgary. All libraries were sequenced on mid-output 150 cycle NextSeq500 runs, with ~130 million reads each. Sequencing (BCL) files were processed with the Cell Ranger 2.1.0 package (10xGenomics). Reads were aligned to the GRCh38 transcriptome. Data standardization and normalization was performed with the Seurat package *(70)*.

### Bulk RNA-seq (Toronto cohort)

RNA from tissue and cells was extracted using Qiagen AllPrep DNA/RNA/miRNA Universal Kit (catalog # 80224). Strand-specific RNA-seq (ssRNA-seq) libraries were constructed from total RNA samples using plate-based protocols. Libraries were sequenced at 75 bp PET using V4 chemistry on a HiSeq 2500 instrument (Illumina) at the Genome Sciences Centre (Vancouver, BC). Reads were aligned with the STAR aligner *(71)* v2.4.2a to hg38 human reference genome (from iGenome). R Bioconductor DESEq2 package *(72)* was used for normalization and vst transformation of the gene expression matrix

### Graphing software

GraphPad Prism 7.0c was used for graph generation. The Venn diagrams were generated using the eulerr R package version 4.0.0 (www.cran.r-project.org/web/packages/eulerr/index.html). Sankey Diagrams were generated using SankeyMATIC (www.sankeymatic.com) and the outputs were modified to be of appropriate format for figures.

### External datasets

Identification of genes included in the OMIM database of genetic inheritance was done with DAVID bioinformatics resources 6.8 *(73, 74)*. Survival analysis of pediatric glioma patients was done with R2: Genomics analysis and visualization platform (www.r2.amc.nl) using a previously published dataset by Paugh et al *(42)*. Assessment of allelic fractions for *ATRX* mutations was done using an independent dataset *(36)* and available at www.pedpancan.com. GTEx data was assessed at https://www.gtexportal.org/home/ *(37)*. The Database of Genomic Variants was accessed at http://dgv.tcag.ca/dgv/app/home *(75)*. Developmental human brain gene expression data was extracted from BrainSpan (http://www.brainspan.org).

### Statistical analysis of enrichment of OMIM genes in the pGBM mutational dataset

The number *X* of genes harboring non-synonymous, stop gain/stop loss, frame shift, non-frame shift SNVs and indels was identified for primary and recurrent tumors. From these, genes with a phenotype in OMIM were selected (*Y*). The OMIM genes (n = 13,520) with a clinical phenotype were identified using the genemap2.txt file (containing gene and phenotype information) downloaded from OMIM (https://www.omim.org/downloads/). For p-value analysis, *X* number of random genes were selected from all protein coding genes in Ensemble (n = 20,345) and among these the number of genes with related OMIM phenotype were selected (Rand_OMIM). This analysis was repeated 100,000 times. The p-value was identified as the number of times “Rand_OMIM’ genes were higher than *Y* number of genes in the 100,000 iterations.

## Acknowledgements

Canadian Institutes of Health Research (CIHR) early career award (Institute of Cancer Research) to MG (ICT-156651); a CIHR project scheme grant to MG (PJT-156278); a Cancer Research Society Scholarship for the Next Generation of Scientists grant to MG; a CIHR graduate scholarship to MH; Royal Society EP Abraham Research Professorship and a Wellcome Trust Senior Investigator Award (098357) to BDS; a Cancer Research Society Scholarship for the Next Generation of Scientists grant to SM. Research supported by SU2C Canada Cancer Stem Cell Dream Team Research Funding (SU2C-AACR-DT-19-15) provided by the Government of Canada through Genome Canada and the Canadian Institute of Health Research, with supplemental support from the Ontario Institute for Cancer Research, through funding provided by the Government of Ontario. Stand Up To Cancer Canada is a Canadian Registered Charity (Reg. # 80550 6730 RR0001). Research Funding is administered by the American Association for Cancer Research International - Canada, the Scientific Partner of SU2C Canada. BDS and DK acknowledge core funding to the Gurdon Institute from the Wellcome Trust (092096) and CRUK (C6946/A14492).

## Authors contribution

MG and MH conceived the experiments; MH, JK, KE and NHD performed the wet lab experiments; MH, AHG, DJK, MJ, AN, KN, MG and SM analyzed the WGS and scRNA-seq data; FMGC, MMK, FJC, BL, YM, AJM, RM, MAM, MDT, GB, TJP and PBD generated the bulk RNA-seq data; JAC, MMK, FJC and PBD provided samples; MG, SM, JAC, BDS and DS provided supervision to the work described hereby; MG, MH and JAC wrote the original manuscript; all authors provided editorial and experimental feedback.

